# Differential induction of interferon stimulated genes between type I and type III interferons is independent of interferon receptor abundance

**DOI:** 10.1101/448357

**Authors:** Kalliopi Pervolaraki, Soheil Rastgou Talemi, Dorothee Albrecht, Felix Bormann, Connor Bamford, Juan Mendoza, Christopher Garcia, John McLauchlan, Thomas Höfer, Megan L. Stanifer, Steeve Boulant

**Author notes:** Corresponding author: (SB). These authors contributed equally to this work (Megan L. Stanifer, Steeve Boulant).

## Abstract

It is currently believed that type I and III interferons (IFNs) have redundant functions. However, the preferential distribution of type III IFN receptor on epithelial cells suggests functional differences at epithelial surfaces. Here, using human intestinal epithelial cells we could show that although both type I and type III IFNs confer an antiviral state to the cells, they do so with distinct kinetics. Type I IFN signaling is characterized by an acute strong induction of interferon stimulated genes (ISGs) and confers fast antiviral protection. On the contrary, the slow acting type III IFN mediated antiviral protection is characterized by a weaker induction of ISGs in a delayed manner compared to type I IFN. Moreover, while transcript profiling revealed that both IFNs induced a similar set of ISGs, their temporal expression strictly depended on the IFNs, thereby leading to unique antiviral environments. Using a combination of data-driven mathematical modeling and experimental validation, we addressed the molecular reason for this differential kinetic of ISG expression. We could demonstrate that these kinetic differences are intrinsic to each signaling pathway and not due to different expression levels of the corresponding IFN receptors. We report that type III IFN is specifically tailored to act in specific cell types not only due to the restriction of its receptor but also by providing target cells with a distinct antiviral environment compared to type I IFN. We propose that this specific environment is key at surfaces that are often challenged with the extracellular environment.

**Author summary:** The human intestinal tract plays two important roles in the body: first it is responsible for nutrient absorption and second it is the primary barrier which protects the human body from the outside environment. This complex tissue is constantly exposed to commensal bacteria and is often exposed to both bacterial and viral pathogens. To protect itself, the gut produces, among others, secreted agents called interferons which help to fight against pathogen attacks. There are several varieties (type I, II, and III) of interferons and our work aims at understanding how type I and III interferon act to protect human intestinal epithelial cells (hIECs) during viral infection. In this study, we confirmed that both interferons can protect hIECs against viral infection but with different kinetics. We determined that type I confer an antiviral state to hIECs faster than type III interferons. We uncovered that these differences were intrinsic to each pathway and not the result of differential abundance of the respective interferon receptors. The results of this study suggest that type III interferon may provide a different antiviral environment to the epithelium target cells which is likely critical for maintaining gut homeostasis. Our findings will also help us to design therapies to aid in controlling and eliminating viral infections of the gut.

## Introduction

During viral infection interferons (IFNs) are the predominant cytokines made to combat viral replication and spread. Following binding to specific receptors, IFNs induce a JAK/STAT signaling cascade which results in the production of interferon stimulated genes (ISGs). These ISGs will then establish an antiviral state within the cell and will also alert surrounding cells and immune cells to assist in viral clearance [1]. There are three classes of IFNs. Type I IFNs are produced by all cell types and are recognized by the ubiquitously expressed heterodimeric receptor IFNAR1/IFNAR2. Type II IFNs are only produced by immune cells [2, 3]. Type III IFNs are made by all cell types but the IFNLR1 (or IL28Ra) subunit of the heterodimeric receptor IFNLR1/IL10RP is restricted to epithelial and barrier surfaces and to a subset of immune cells [4-9]. Despite the fact that type I and type III IFNs are structurally unrelated and engage different receptors, signaling downstream of both receptors exhibits a remarkable overlap and leads to the induction of a similar pool of ISGs. These observations originally led to the hypothesis that type I and III IFNs were functionally redundant.

This model has been challenged more and more in recent studies which highlight that the cell type specific compartmentalization of IFNLR1 provides type III IFNs a unique potential for targeting local infections at mucosal surfaces. For example, *in vivo* data on enteric virus infection of the murine gastrointestinal tract showed that responsiveness to type III IFN is necessary and sufficient to protect murine intestinal epithelial cells (IECs) against rotavirus and reovirus infection [10-12]. On the contrary, type I IFN was necessary to protect against viral infection of cells in the lamina propria and against systemic spread [10, 11]. Likewise, it was demonstrated that fecal shedding of norovirus was increased in IFNLR1-deficient, but not IFNAR1-deficient, mice, showing that type III IFN uniquely controls local norovirus infection in the gut [13, 14]. Similarly, in the respiratory tract, type III IFNs are predominately produced upon infection with influenza A virus [15-19]. However, as infection progresses type I IFN comes into play to reinforce viral inhibition by inducing a pro-inflammatory response [20].

Differences in the antiviral activity conferred by both cytokines appear to be not only driven by the spatial restriction of their receptors but also by intrinsic subtle differences in signal transduction. It was demonstrated, in human hepatocytes and lung epithelial cells, that type I IFN confers a more potent antiviral protection compared to type III IFNs [5, 21-23]. Additionally, it was shown in human IECs that type III IFN partially requires MAP kinase activation to promote an antiviral state while type I IFN was independent of it [24]. Although it has been reported in many studies that very similar ISGs are induced upon type I or type III IFN stimulation of cells, work mostly performed in hepatocytes revealed that both cytokines induce these ISGs with different kinetics [21, 25-27]. Type III IFN mediated signaling was found to be associated with a delayed and reduced induction of ISGs compared to type I IFNs [25, 26]. Similar differences in the magnitude and/or kinetics of ISGs induction upon type I versus type III IFN treatment were observed in human primary keratinocytes, airway epithelial cells and in Burkitt’s lymphoma derived B (Raji) cells, as well as in murine intestinal and lung epithelial cells and immune cells [20, 28-31].

The molecular mechanisms leading to this delayed and reduced induction of ISGs upon type III IFN treatment remains unclear. As these differences in kinetics of ISG expression between both IFNs could not be directly explained by their signaling cascades an alternative explanation was proposed where type III IFN receptor is expressed at lower levels at the cell surface. This lower receptor expression level could provide a biochemical explanation for the observed differences in delayed kinetics and weaker amplitude of ISG expression compared to type I IFN. However, to date, there is no direct experimental evidence for this model. Similarly, whether the observed differences between both IFNs is intrinsic to both specific signal transduction pathways and whether it is restricted to some cell types (e.g. hepatocytes) or represents a global signaling signature in all cells expressing both IFN receptors has not been fully addressed.

In this study, we have investigated how type I and III IFNs establish their antiviral program in human mini-gut organoids and human IEC lines. We found that type I IFN can protect human IECs against viral infection faster than its type III IFN counterpart. Correspondingly, we determined that type I IFN displays both a greater magnitude and faster kinetics of ISG induction compared to the milder, slower type III IFN. By developing mathematical models describing both type I and type III IFN mediated production of ISGs and by using functional receptor overexpression approaches, we demonstrated that the observed lower magnitude of ISG expression for type III IFNs was partially the result of its lower receptor expression level compared to the type I IFN receptor. Inversely, the observed delayed kinetics of type III IFN cannot be explained by receptor expression level indicating that this property is specific to type III IFN and inherent to its signaling pathway. Our results highlight important differences existing between both type I and type III IFN-mediated antiviral activity and ISG expression which are not only the result of receptor compartmentalization but also through intrinsic fundamental differences in each IFN-mediated signaling pathway.

## Results

### Type III IFN-mediated antiviral protection is delayed compared to type I IFN

We have previously reported that both type I and III IFNs mediate antiviral protection in human IECs [24]. To address whether type I and type III IFN have a different profile of antiviral activity in primary non-transformed human IECs, as reported in human lung cells [22], we compared the antiviral potency of both IFNs in human mini-gut organoids. Colon organoids were pretreated with increasing concentrations of either type I or III IFNs for 2.5 hours and subsequently infected with vesicular stomatitis virus expressing luciferase (VSV-Luc). Viral infection was assayed by bioluminescence and results showed that both IFNs induced an antiviral state in a dose-dependent manner. We observed that type I IFN was slightly more potent in protecting against viral infection at higher concentration compare to type III IFNs. Type I IFN could almost fully inhibit viral infection while type III IFN was only able to reduce infection to around 80% (Fig 1A). Interestingly, the concentration of type I IFN necessary to provide 90% of relative antiviral protection (EC90) was significantly lower than the one for type III IFN (Fig 1B).

**Fig.1.**
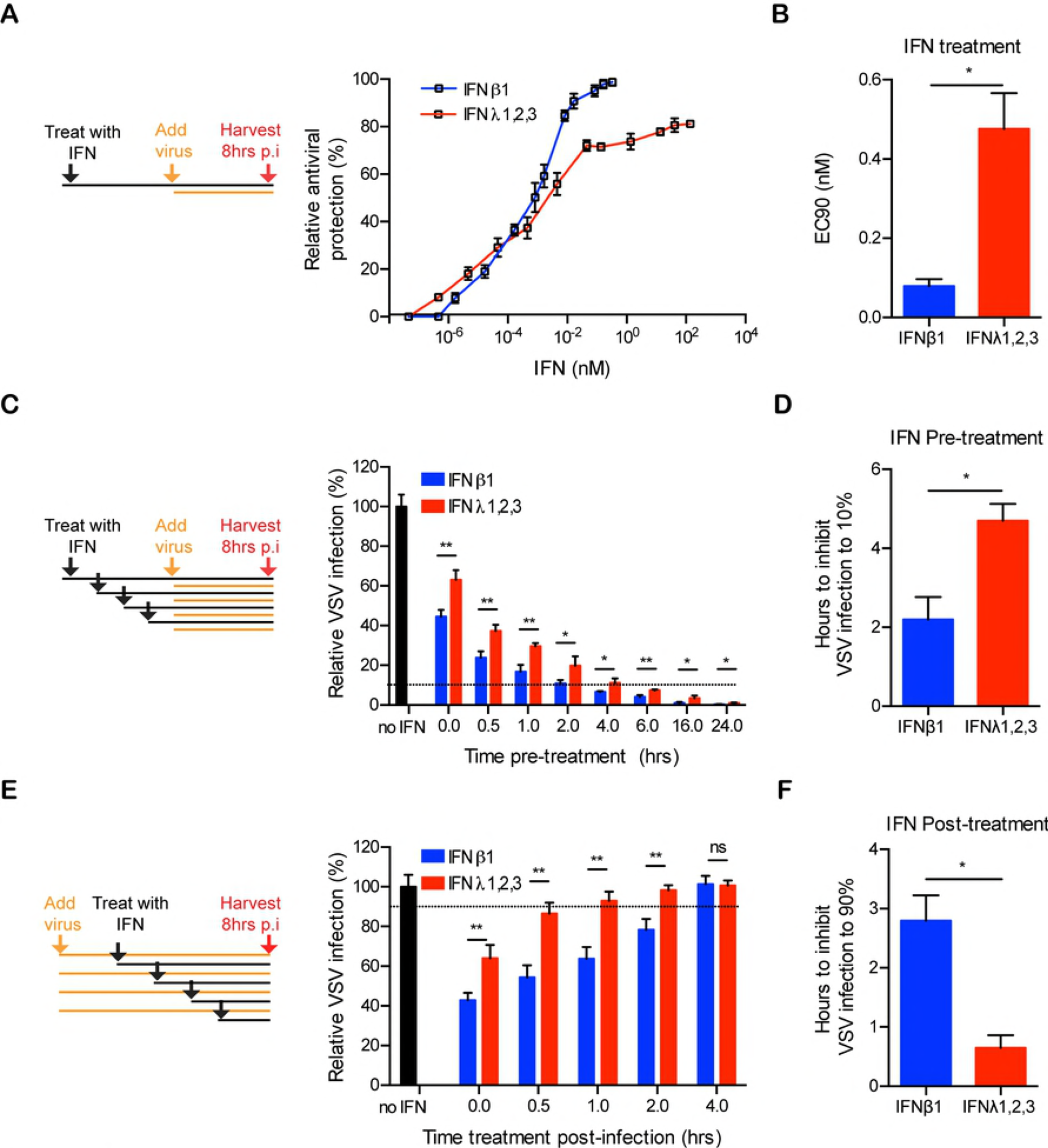
Kinetics of type I and type III IFN-mediated antiviral activities in human mini gut-organoids. (A-B) Colon organoids were pre-treated with the indicated concentrations of type I IFN (P) or type III IFN (A1-3) for 2.5 h prior to infection with vesicular stomatitis virus (VSV) expressing Firefly luciferase (VSV-Luc) using a multiplicity of infection (MOI) of 1. Viral replication was assayed by measuring the luciferase activity. (A) The relative antiviral protection is expressed as a percentage of total protection in VSV-infected organoids or (B) as the EC90 corresponding to the concentration of type I IFN (P) or type III IFN (A1-3) resulting in 90% inhibition (10% infection) of viral replication. (C-D) Colon organoids were treated with type I IFN (P) (2,000 RU/mL equivalent 0.33 nM) or type III IFN (A1-3) (100ng/mL each or total 300 ng/mL equivalent 13.7 nM) for different times prior to infection with VSV-Luc. Viral replication was assayed by measuring luciferase activity. (C) The relative VSV infection is expressed as the percentage of the luciferase activity present in VSV-infected organoids without IFN treatment (set to 100). (D) Preincubation time of type I IFN (P) or type III IFN (A1-3) required to inhibit VSV infection to 10% (90% inhibition). (E-F) Same as (C-D), except colon organoids were treated at the indicated times post-infection with VSV-Luc. (F) Delayed-time post-infection for type I IFN (P) or type III IFN (A1-3) to still inhibit VSV infection to 90% (10% inhibition). Data in (A-F) represent the mean values of two independent experiments. Error bars indicate the SD. *<P.05, **P < 0.01, ns, not significant (unpaired t-test).

To determine whether type III IFN requires a prolonged treatment to achieve similar antiviral protection as observed with type I IFN, we performed a time course experiment in which human colon organoids were pre-treated for different times with either IFN prior infection with VSV-Luc (Fig 1C). We found that approximately 2 hours pre-treatment with type I IFN was sufficient to reduce VSV infection by 90% (10% remaining infection), while type III IFN required around 5 hours to achieve a 90% reduction of infectivity (Fig 1C and 1D). Interestingly, 24 hours of pretreatment was necessary for type III IFN to almost completely prevent VSV infection (Fig 1C). These results strongly suggest that both type I and type III IFN could have similar potency but that type III IFN requires more time to establish an antiviral state.

We next addressed how long after infection IFN treatment is still able to promote antiviral protection. Colon organoids were infected with VSV-Luc and treated at different times post-infection with either type I or III IFNs. Interestingly, type I IFN could inhibit viral replication even when added several hours post-infection. In contrast, type III IFN appeared to require a much longer time to establish its antiviral activity and was unable to efficiently protect the organoids after VSV infection has initiated (Fig 1E and 1F).

Importantly, these differences in the kinetics of antiviral activity of type I versus type III IFNs were neither donor nor colon specific as similar results were observed in intestinal ileum-derived organoids derived from different donors (Sup Fig 1). In addition, human colon carcinoma-derived cell lines T84 (Sup Fig 2) and SKCO15 (data not shown) fully phenocopy the difference in type I versus type III IFN antiviral activity generated by primary mini-gut organoids. Taken together these results demonstrate that while both type I and III IFNs can promote similar antiviral states into target cells, they do so with distinct kinetics. The cytokine-induced antiviral state is promoted faster by type I IFN compared to type III IFN.

### Type I and III IFNs induce different amplitudes and kinetics of ISG expression

To understand how type I and type III IFNs promote an antiviral state in primary IECs but with different kinetics, we analyzed the magnitude of ISG expression over time upon IFN treatment. Colon organoids were treated with increasing concentrations of either type I or type III IFN and the expression levels of two representative ISGs (IFIT1 and Viperin) were assayed at different times post-IFN treatment. Results revealed that type I IFN ultimately leads to a significantly higher induction of both IFIT1 and Viperin compared to type III IFN (Fig 2A and 2B). This difference in the magnitude of ISG stimulation was independent of the duration of IFN treatment (Fig 2A and 2B). To determine if this pattern of expression applies to other ISGs, we treated colon organoids with either type I or type III IFN over a 24-hour time course, and analyzed the mRNA levels of 132 different ISGs and transcription factors involved in IFN signaling (see complete list of genes and corresponding primers in Sup Table 1 and 2) (Fig 2C and 2D). Differential expression analysis revealed that both type I and type III IFNs induce almost the same set of ISGs and that most of the genes significantly induced by type III IFN were also induced by type I IFN (Fig 2C). However, similar to IFIT1 and Viperin (Fig 2A and 2B), we found that the magnitude of ISG expression was greater for type I IFN compared to type III IFN (Fig 2D). Similar results were found in the immortalized colon carcinoma-derived T84 cells (Sup Fig 3A-C).

**Fig.2.**
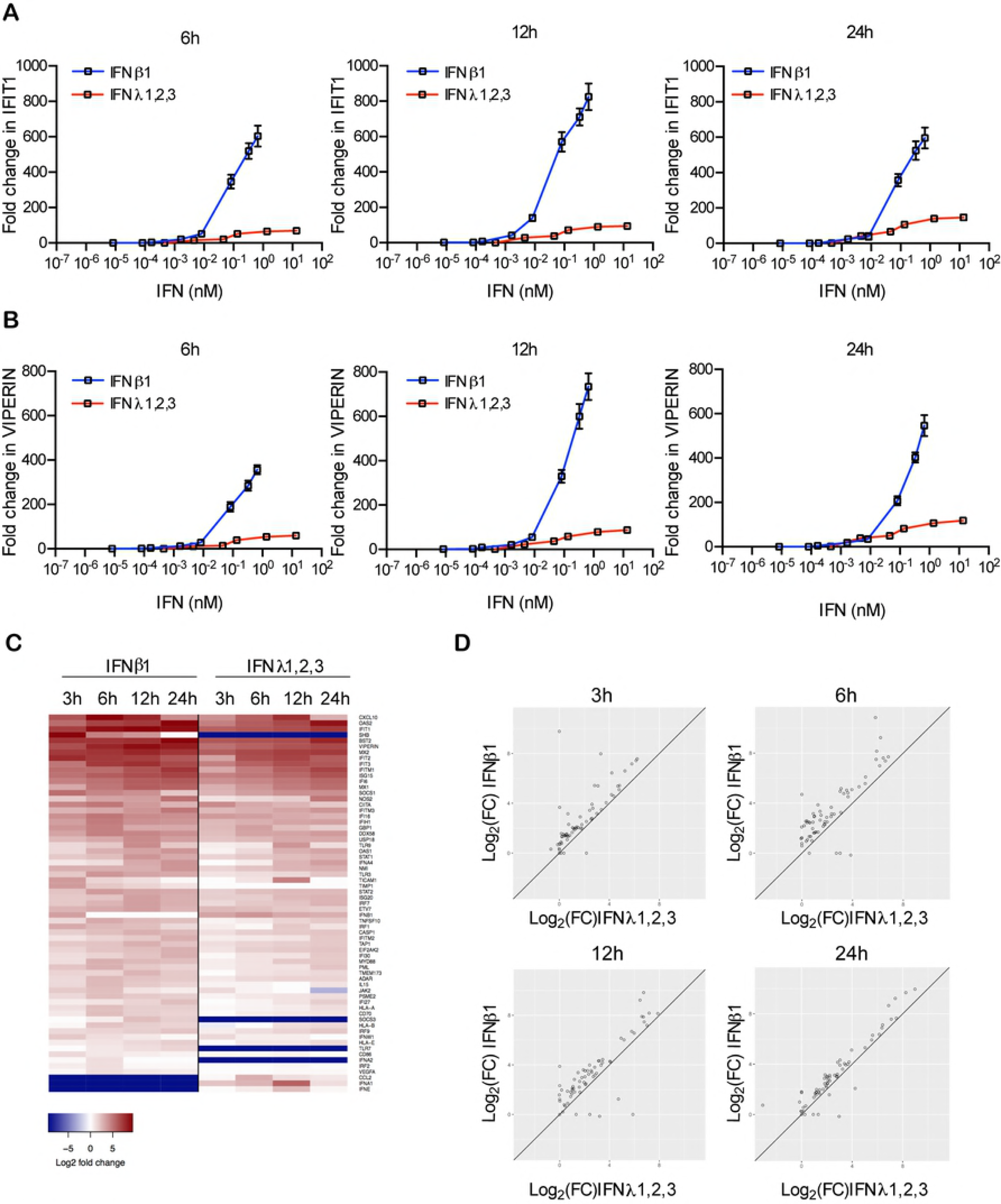
Type III IFNs have a lower transcriptional activity compared to type I IFNs. (A-B) Human colon organoids were stimulated with indicated concentrations of type I (P) or III IFN (A1-3) for different times and the transcript levels of the ISGs IFIT1 and Viperin were analyzed by qRT-PCR. Data are normalized to TBP and HPRT1 and are expressed relative to untreated samples at each time point. A representative experiment with technical triplicates, out of three independent experiments is shown. Mean values and SD are shown. (C) Colon organoids were treated with type I IFN (P) (2,000 RU/mL equivalent 0.33 nM) or type III IFN (A1-3) (300 ng/mL equivalent 13.7 nM) for the indicated times and identification of the IFN-induced ISGs was performed by qRT-PCR. A total of 65 out of 132 ISGs tested were found to be significantly induced more than 2-fold compared with a baseline (mean of untreated controls at the particular time points) for at least one time point by at least one IFN treatment. Data are normalized to TBP and HPRT1 and visualized in a heatmap using R after sorting the fold change of expression in response to type I IFN (P) in decreasing order. (D) Comparison of expression values (log2 (Fold Change)) for all genes induced at the indicated times with type I IFN (P) versus type III IFN (A1-3). Solid line indicates equivalent expression.

To address whether there is any correlation between the different antiviral protection kinetics conferred by type I and III IFNs (Fig 1) and the kinetics of ISG expression, we analyzed the temporal expression of ISGs upon IFN treatment of human colon organoids. Hierarchical clustering analysis of all ISGs up-regulated upon type I or type III IFN treatment defined four distinct expression profiles based on the time of their maximum induction (Fig 3A-C). Group 1 are ISGs whose expression peaks 3 hours post-IFN treatment. The expression of ISGs in group 2 and 3 peaks at 6 and 12 hours post-treatment, respectively. Group 4 corresponds to ISGs with a continuous increase in expression over time (Fig 3A and B). Under type I IFN treatment, ISGs were nearly equally distributed in all four expression groups (Fig 3A, 3C, and 3D). By contrast, although the same ISGs were induced by type III IFN, they almost all belong to the expression group 4, being expressed later after IFN treatment (Fig 3B-D). In line with the primary mini-gut organoids, T84 cells presented similar differences in the kinetics of ISGs expression (Sup Fig 3D). Importantly, cell polarization of human IECs did not impact the kinetics of ISG expression as similar results were obtained when comparing polarized vs. non-polarized T84 cells (data not shown).

**Fig.3.**
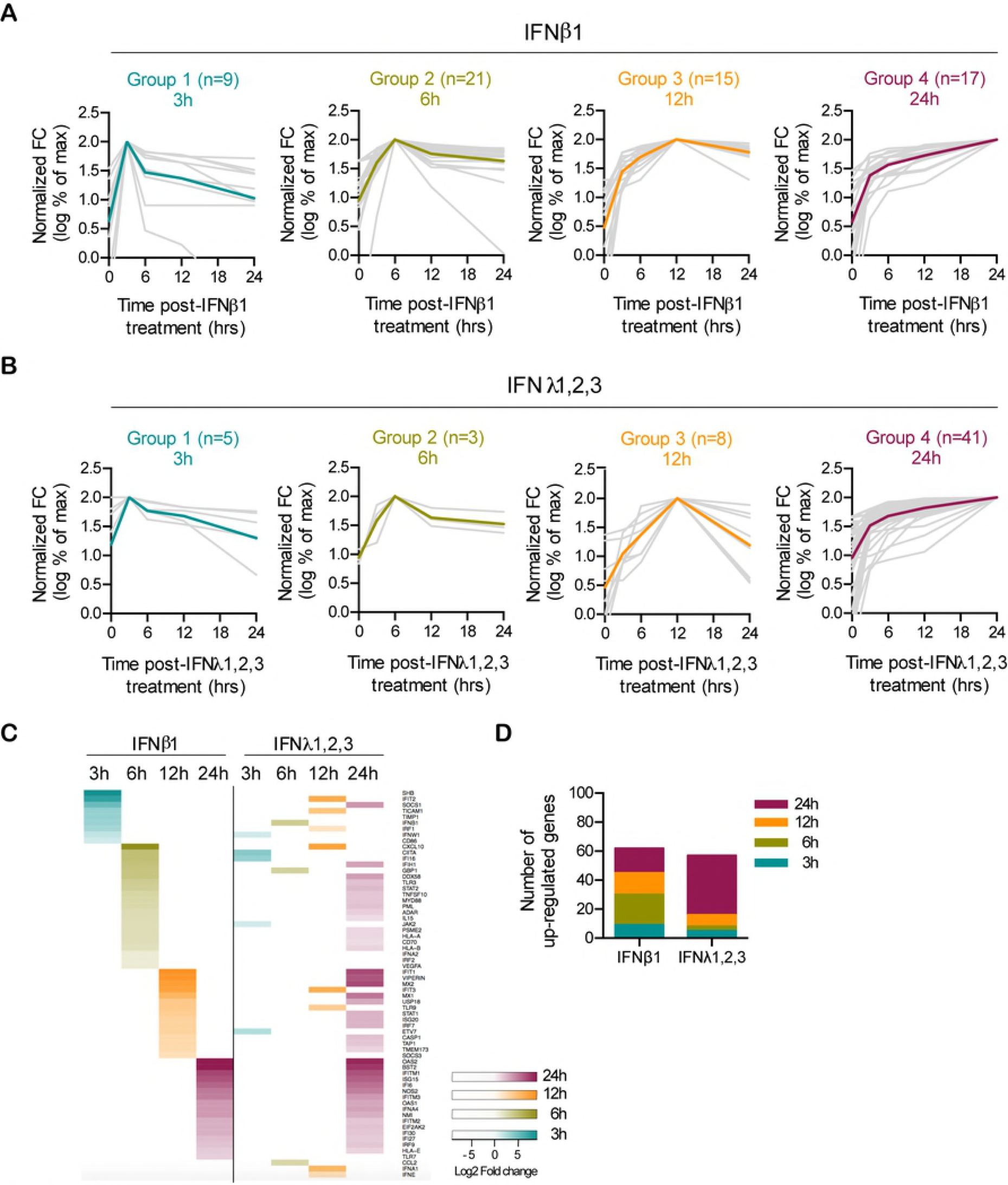
Type III IFNs present delayed transcriptional activity compared to type I IFNs. (A-D) Human colon organoids were treated with type I IFN (P) (2,000 RU/mL equivalent 0.33 nM) or type III IFN (A1-3) (300 ng/mL equivalent 13.7 nM) for 3, 6, 12 or 24 hours and the kinetic pattern of expression of the 65 significantly up-regulated ISGs were analyzed by qRT-PCR in triplicates. Data are normalized to TBP and HPRT1 and are expressed relative to untreated cells at each time point. Hierarchical clustering analysis of these genes produced four distinct temporal expression patterns (Groups 1-4) based on the time-point of the maximum induction in response to type I IFN (P) or type III IFN (A1-3). Color codes have been used to visualize the induction peak per group. (A-B) Gray lines show the normalized kinetic expression of each gene for each group upon treatment with (A) type I IFN (P) or (B) type III IFN (A1-3). The colored lines are the average of the kinetic profiles for the genes of each group. (C) Gene expression heat map showing the genes clustered in their respective temporal expression patterns groups in response to type I IFN (P) or type III IFN (A1-3). The genes per group are sorted in decreasing order on the basis of their fold change of expression in response to type I IFN (P) or type III IFN (A1-3) and only showing the highest expressed values within the temporal groups omitting all other values for visualization. (D) Number of genes belonging to each group.

We next wanted to control that our observed differences in the kinetics of ISGs expression induced by both cytokines were independent of IFN concentration. Colon organoids were treated with increasing amounts of type I or type III IFNs and the transcriptional up-regulation of representative ISGs belonging to each of the expression profile groups (group 1-4) was measured over time (Fig 4). Consistent with our previous results, the temporal expression patterns of each representative ISGs were independent of the IFN concentration and the ISG expression kinetic signature was specific to each IFN (Fig 4). Complementarily, to address whether the observed differences between type I and type III IFNs were not due to the lower affinity of type III IFN for its receptor compared to type I IFN, we employed the high affinity variant of type III IFN (H11-IFNλ3) [32] to monitor the kinetics of ISG expression. Results show that cells treated with the higher affinity H11-IFNλ3 display a higher magnitude of ISG expression but their kinetics of expression were unchanged (Sup Fig 4). Altogether, our results strongly suggest that although both type I and type III IFNs induce a similar set of ISGs in hIECs, type III IFN induces globally a lower amplitude and a delayed ISG expression compared to type I IFN.

**Fig.4.**
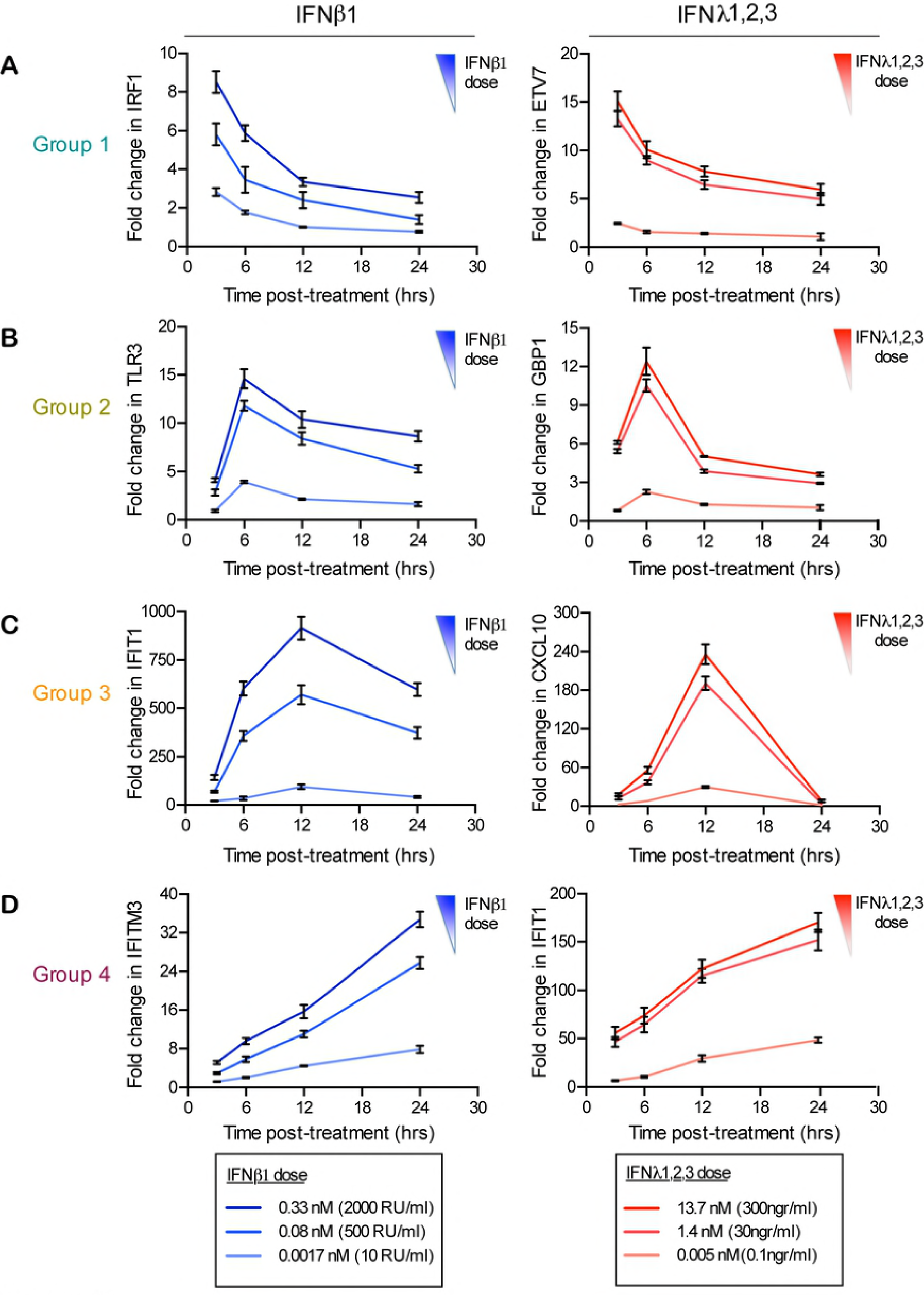
Validation of the unique kinetic patterns of ISG expression upon type I versus type III IFN treatment. (A-D) Human colon organoids were stimulated with increasing concentrations of type I IFN (P) or III IFN (A1-3) for indicated times and the kinetic pattern of expression of one representative ISG from each temporal expression patterns groups 1-4 was analyzed by qRT-PCR, (left column) type I IFN (P), (right column) type III IFN (A1-3) treated organoids. Data are normalized to HPRT1 and are expressed relative to untreated cells at each time point. A representative experiment with technical triplicates. Mean values and SD are shown.

### Mathematical modeling shows that IFN receptor abundance modulates the magnitude of ISG response while the type I and type III IFN specific kinetic profiles are independent of receptor abundance

Our data show remarkable differences in the magnitude and kinetics of ISGs induced by type I versus type III IFN (Fig 2-3 and Sup Fig 3), and in the subsequent induction of a differential antiviral state (Fig 1 and Sup Fig 1-2). To investigate the mechanisms underlying these differences, we used data-driven mathematical modeling and model selection. We considered three mechanistic causes for the observed differential signaling: (1) Receptor abundance (different number of IFNLR compared to IFNAR complexes); (2) Receptor regulation (different rates of activation and/or inactivation of IFNLR compared to IFNAR complexes); (3) STAT activation (different rates of STAT activation by type I and type III IFNs). We devised corresponding mathematical models describing the dynamics of receptor activation and inactivation, STAT1/2 phosphorylation and STAT-dependent activation of ISG expression (Fig 5A). The models were implemented as systems of ordinary differential equations (Sup Table 3) and fitted to the time-resolved data for the prototypical ISG, Viperin, measured with different doses of type I or type III IFNs and with the high affinity H11-IFNλ3. We ranked the models according to the Akaike information criterion corrected for small sample size (AICc), which evaluates the goodness of fit and, at the same time, penalizes the number of fit parameters (for more details see Materials and Methods). Throughout, we allowed different receptor abundance, but this difference alone could not account for the different signaling dynamics (Fig 5B; model Mi has negligible support by the data, as quantified by the small AICc weight, which is a weight of evidence for the respective model). Interestingly, in addition to receptor abundance, the best-fitting model (M_3_) has also different rates of activation and inactivation of IFNLR and IFNAR complexes. However, alternative models with different rates of STAT activation and/or ISG expression have good performance (M_2_ and M_4_, respectively). Therefore, the modeling indicates that differential ISG activation by type I and type III IFNs is likely due to different abundance of the respective receptors and cell-intrinsic differences in how the signals from bound receptors are processed.

**Fig.5.**
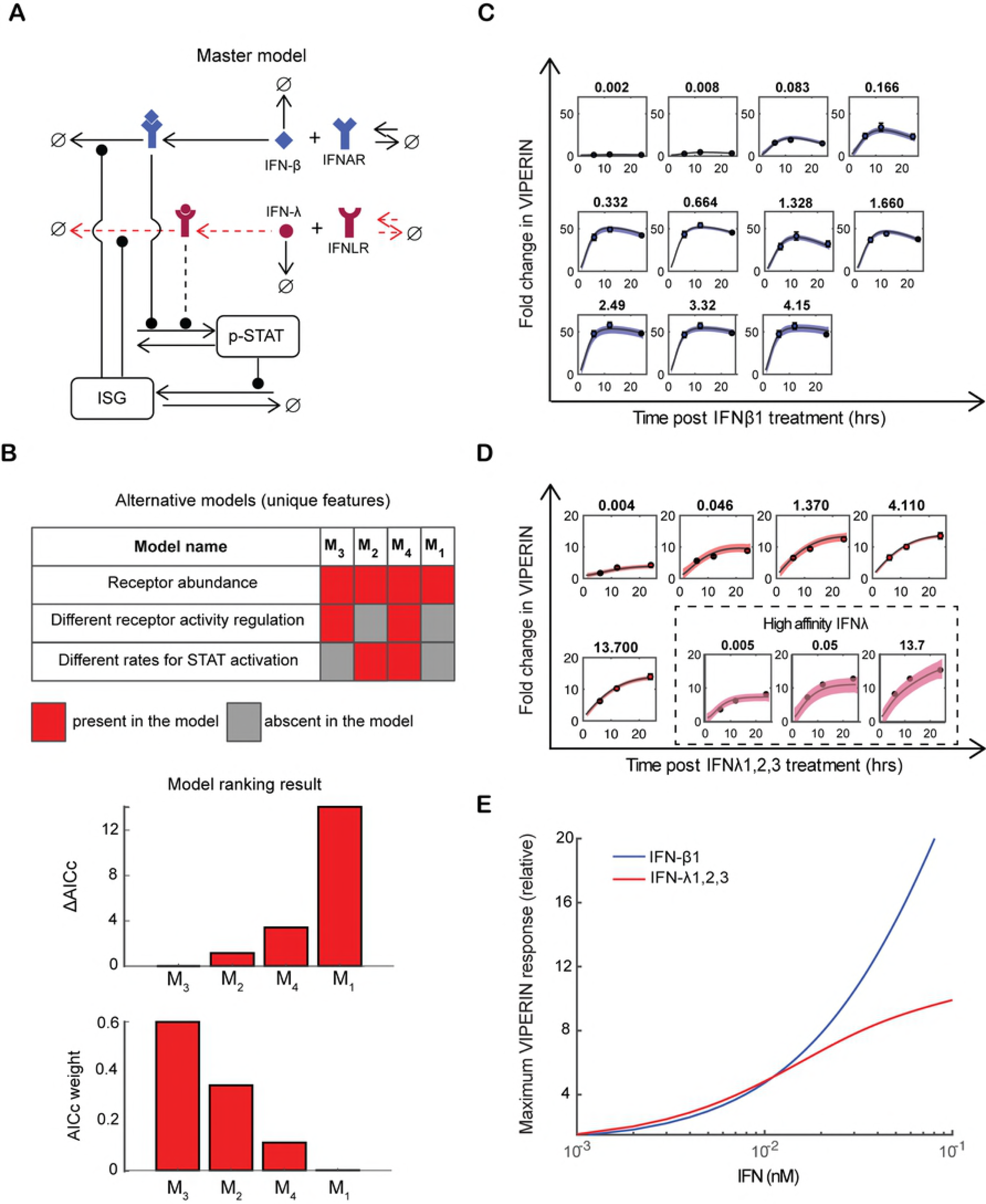
Mathematical modeling of type I and type III IFN responses. (A) Scheme of the mathematical model. IFNs bind to their cognate receptors and activate them; all molecules are also subject to degradation (0). Active receptors mediate STAT phosphorylation while phosphorylated STAT (p-STAT) drives ISG expression. ISGs may include negative feedback regulator of STAT activation. Dashed lines indicate the potential sources of difference between the two pathways. Red dashed lines show the sources of the difference between the two pathways implemented in the best-fitting model. (B) Model selection. Models fitted to the experimental data were ranked using the Akaike information criterion corrected for small sample size (AlCc) and the AlCc weight, as a measure of support for the given model by the data. (C-D) The best-fitting model M3 reproduces the Viperin expression dynamics upon treatment with different concentrations of (C) type I IFN and (D) type III IFN (see Sup Fig 3A-B for experimental data). In (C) and (D), the solid lines represent the best fits and the shaded areas are 95% confidence intervals. (E) Simulation of the maximum Viperin induction upon treatment with equal concentrations of type I IFN or type III IFN.

The best-fitting model (M3) accounted for the dose-response and the different Viperin expression kinetics triggered by type I, type III and the high affinity H11-IFNA3 in T84 cells, group 3 and group 4 expression kinetics, respectively (Fig 5C, D). The different kinetics of the IFN responses-fast and transient for type I IFN vs slower and sustained for type III IFN-are predicted to be largely due to receptor inactivation, which is faster for IFNAR than for IFNLR complex (Sup Fig 5A-C). Interestingly, the model shows that at low IFN concentrations, Viperin is induced almost equally by both IFNs whereas at higher concentrations, type I IFN induces Viperin more strongly (Fig 5E). These dose-dependent features agree with our experimental data (Sup Fig 3B, right panel).

Next, we tested the pivotal impact of receptor expression on ISG induction that was indicated by our model. Specifically, the model predicts that an increase in IFNAR1 or IFNLR1 level will increase the amplitude of ISG induction while preserving the specific kinetic profiles elicited by the two types of IFNs (Sup Fig 5D-E). To experimentally validate the model predictions, IFNAR1 and IFNLR1 were overexpressed in T84 cells. Overexpression of the respective IFN receptor chain was confirmed by reverse quantitative PCR (Sup Fig 6). To ensure the functionality of both IFN receptors, IFNAR1 or IFNLR1 were expressed in our previously characterized knockout T84 cell lines deficient for either the IFN alpha receptor 1 (IFNAR1-/-) or the IFN lambda receptor 1 (IFNLR1-/-) (Sup Fig 7A and 7E) [24]. Our results show that overexpression of IFNAR1 in our IFNAR1-/-T84 cells (IFNAR1-/-+rIFNAR1) restores their antiviral activity, their ability to phosphorylate STAT1 and induce the production of the ISGs IFIT1 and Viperin in the presence of type I IFN (Sup Fig 7B-D). Similarly, although IFNLR1-/-cells were insensitive to type III IFN treatment, overexpression of IFNLR1 (IFNLR1-/-+rIFNLR1) restored their antiviral activity, pSTAT1 and ISG induction after addition of type III IFN (Sup Fig 7F-H). These results demonstrate the functionality of both IFN receptors and validate our overexpression approach as a means to increase IFNAR1 and IFNLR1 levels at the cell surface.

Wild-type T84 cells overexpressing type I IFN receptor (WT+rIFNAR1) were treated with increasing concentrations of type I IFN. Our results showed elevated levels of STAT1 phosphorylation and ISG induction in response to stimulation with type I IFN compared to wild-type cells (Fig 6A-D). Importantly, the response of T84 cells overexpressing type I IFN receptor to type III IFN remained unchanged, indicating a selective enhancement of the type I IFN signaling pathway. Similarly, overexpression of type III IFN receptor (WT+rIFNLR1) shows a significant increase in phosphorylated STAT1 and ISG expression compared to wild-type cells upon type III IFN stimulation, while no difference was observed upon type I IFN treatment (Fig 6E-H). Altogether, our experimental data are consistent with the modeling predictions and confirm the crucial impact of surface receptor levels for regulating the magnitude of type I and III IFN response.

**Fig.6.**
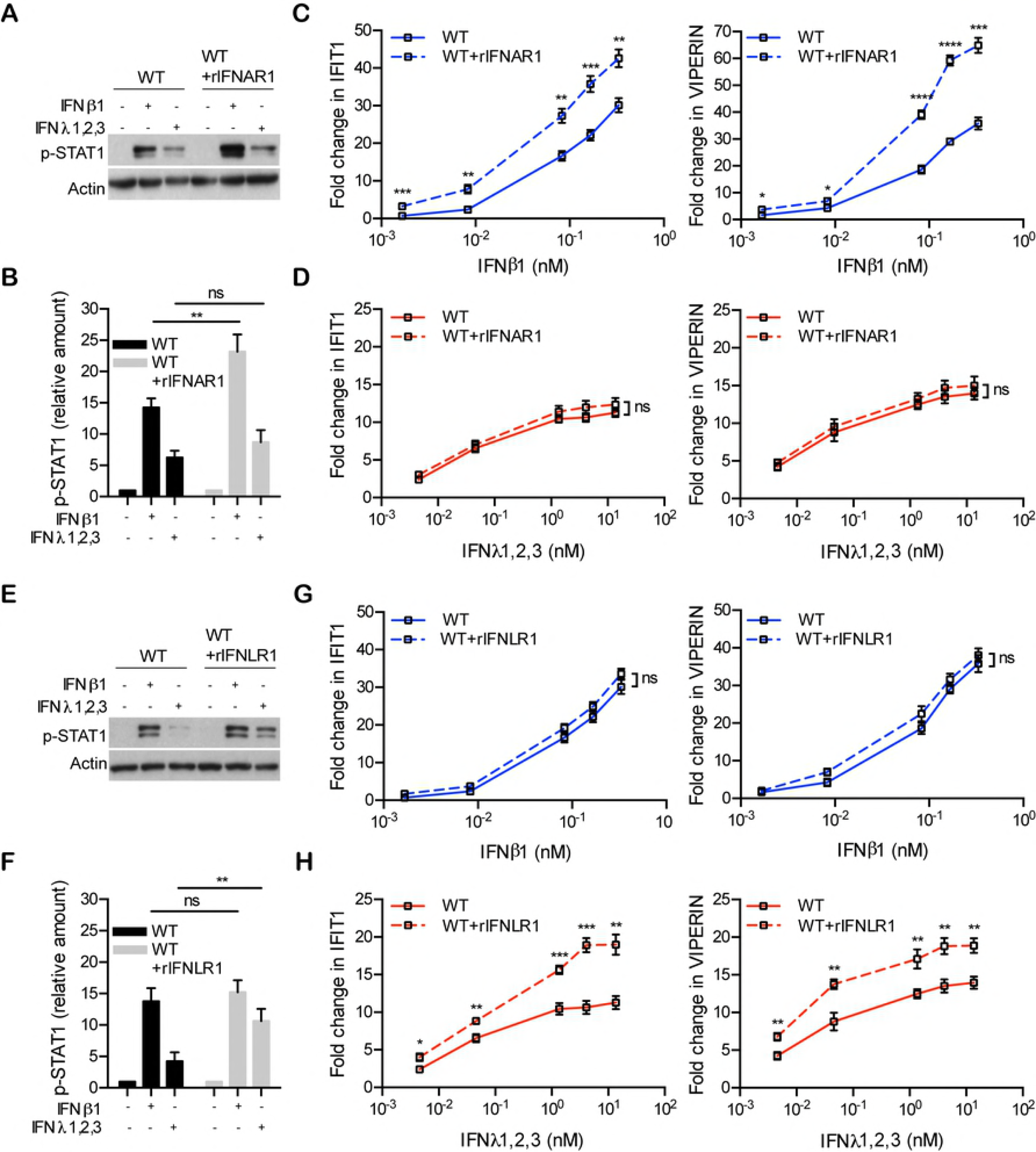
Overexpression of type I and type III IFN receptor increases the transcriptional activity of both cytokines. (A-F) Wild-type T84 cells were transduced with rIFNAR1 or rIFNLR1 to create stable lines overexpressing either IFN receptors. (A-B) T84 wild-type cells (WT) and T84 cells overexpressing rIFNAR1 (WT+rIFNAR1) were treated with type I IFN (P) (2,000 RU/mL equivalent 0.33 nM) or type III IFN (A1-3) (300 ng/mL equivalent 13.7 nM) for 1h and IFN signaling was measured by immunoblotting for pSTAT1 Y701. Actin was used as a loading control. A representative immunoblot out of three independent experiments is shown. (C-D) T84 wild type cells (WT) and T84 cells overexpressing rIFNAR1 (WT+rIFNAR1) were treated with increasing concentrations of type I IFN (P) for 12 hours or type III IFN (A1-3) for 24 hours and the transcript levels of the ISGs IFIT1 and VIPERIN were analyzed by qRT-PCR. Data are normalized to HPRT1 and are expressed relative to untreated cells at each time point. (E-H) Same as (A-D), except T84 cells overexpressing rIFNLR1 (WT+IFNLR1) were used. The mean value obtained from three independent experiments is shown. Error bars indicate the SD. *<P.05, **P < 0.01, ***P < 0.001, ****P < 0.0001, ns, not significant (unpaired t-test).

We next addressed whether this increase of ISG expression in cells overexpressing either the type I or type III IFN receptor was associated with an improved antiviral activity. Wild-type T84 cells overexpressing type I IFN receptor (WT+rIFNAR1) were treated with type I IFN at different time points prior to infection with VSV-Luc virus and their antiviral activity was compared to wild-type T84 cells. Our results showed that the potency and the kinetics of the antiviral activity of cells overexpressing type I IFN receptor does not present any significant change upon type I IFN treatment (Sup Fig 8A). Similarly, there is no difference in the antiviral activity when cells overexpressing type I IFN receptor were treated with type I IFN at different time points post-infection (Sup Fig8B). However, overexpression of type III IFN receptor (WT+rIFNLR1) shows a modest but significant enhancement in type III IFN antiviral potency in the earlier time points of pre-treatment (between 30 minutes and 2 hours) compared to wild-type cells upon type III IFN stimulation (Sup Fig 8G), while they responded similarly to wild-type cells upon type I IFN treatment (Sup Fig 8E). Consistent with this, cells overexpressing type III IFN receptor are more protected than wild-type cells when type III IFN was added post-infection for the early time points (Sup Fig 8H).

Finally, to experimentally validate the limited impact of the IFN receptors abundance on the kinetic profile of ISG expression, as predicted by the model (Sup Fig 5D-E), wild-type cells overexpressing either of the IFN receptors were treated with increasing doses of type I or type III IFNs and the expression of a representative ISG belonging to each of the expression profile groups (group 1-4, Fig 3) was analyzed over time (Fig 7A-D). The experimental data show that the amplitude of ISG expression was dependent on both the dose of IFNs used to stimulate the cells and on the expression levels of the IFN receptors (Fig 7A-D). Importantly, the kinetic profile of ISG expression was similar between WT cells and cells overexpressing the IFNAR1 (WT+rIFNAR1), independent of the applied IFN type I dose (Fig 7A-D left panel). Similarly, wild-type cells overexpressing the IFNLR1 (WT+rIFNLR1) showed no change in the kinetic profile of ISG induction upon type III IFN stimulation (Fig 7A-D right panel). Moreover, we found that the model reproduced the kinetic dose-response data when the IFNAR1 and IFNLR1 expression levels were increased ~2.6 and ~1.5 times, respectively, while all other parameters were held constant (Sup Fig 9). Indeed, we found that IFNAR1 overexpression was stronger than IFNLR1 overexpression, as judged by the transcript levels (Sup Fig 6B-C), with the ratio being consistent with the model prediction (Sup Fig 9D and Sup Fig 6B-C).

**Fig.7.**
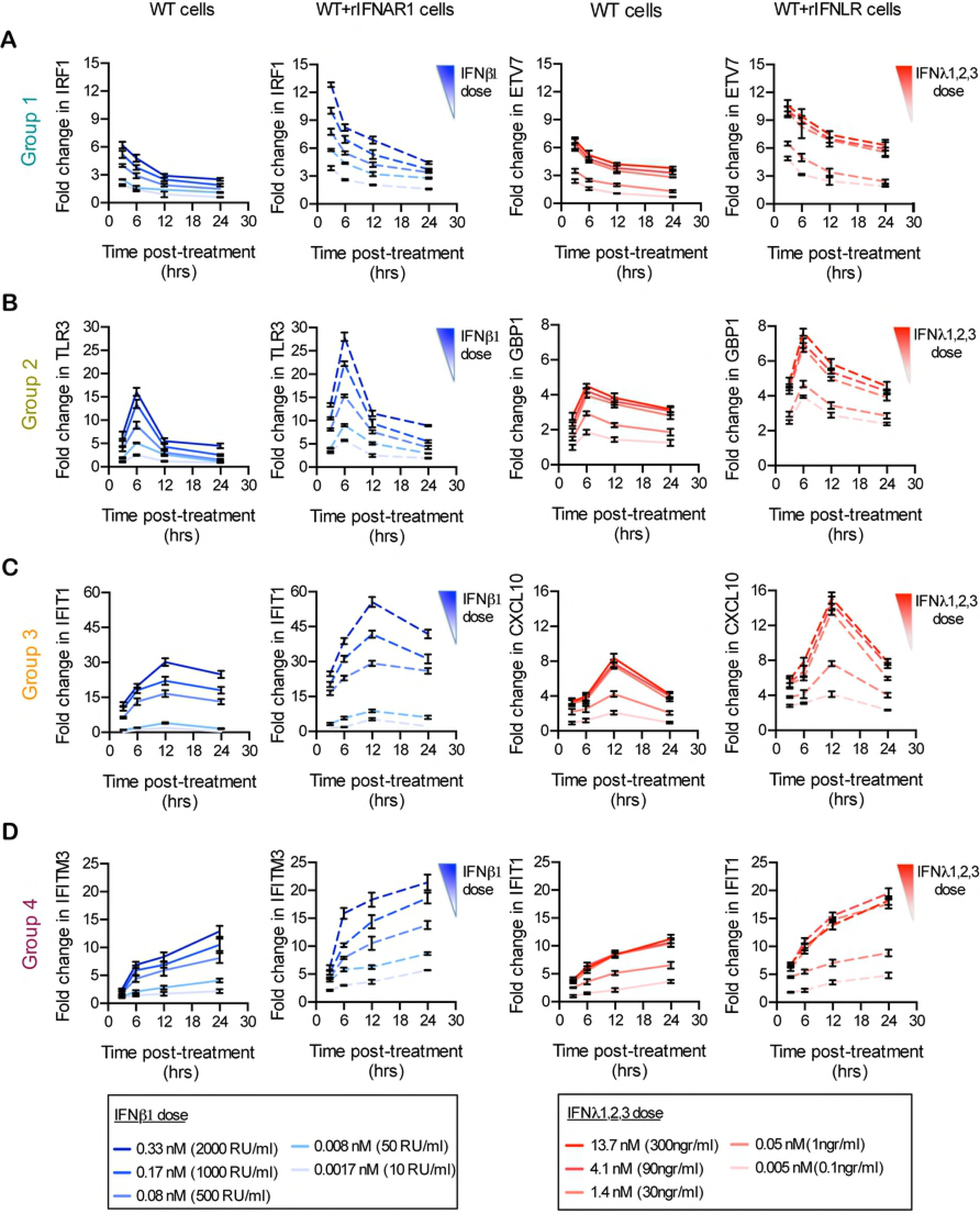
Expression kinetics of ISGs are independent of the IFN receptor levels. (A-D) Wild-type T84 cells were transduced with rIFNAR1 or rIFNLR1 to create stable lines overexpressing either receptors. (blue panels) T84 wild-type cells (WT) and T84 cells overexpressing the IFNAR1 (WT+rIFNAR1) were treated with increasing concentrations of type I IFN (P) for the indicated times and the kinetic pattern of expression of one representative ISG from each temporal expression patterns groups 1-4 was analyzed by qRT-PCR. Data are normalized to HPRT1 and are expressed relative to untreated cells at each time point. (red panels) Same as (blue panels), except T84 cells overexpressing the IFNLR1 (WT+IFNLR1) were used and treated with increasing concentrations of type III IFN (A1-3). A representative experiment with technical triplicates, out of three independent experiments is shown. Mean values and SD are shown.

To directly correlate ISG expression kinetics and amplitude with the expression level of the type III IFN receptor, we thought of overexpressing an IFNLR1 tagged with the GFP fluorescent protein (IFNLR-GFP) in human IECs. To control the functionality of the GFP tagged receptor, the IFNLR1-GFP construct was overexpressed in the human embryonic kidney cell line 293 HEK, which normally elicit a very limited response upon type III IFN treatment. Quantitative RT-PCR revealed that 293 HEK cells overexpressing IFNLR1-GFP produced significantly more ISGs upon type III IFN treatment compared to WT 293 HEK cells or 293 HEK cells expression GFP alone (data not shown). Wild-type T84 cells overexpressing the IFNLR1-GFP (WT+rIFNLR1-GFP) were treated with type III IFN over time and cells were sorted by flow cytometry based on their level of IFNLR1-GFP expression (no GFP expressing (neg), or low and high GFP expressing cells) (Fig 8A). The induction of a representative ISG belonging to each of the expression profile groups (group 1-4, Fig 3) was measured over time in each sorted population (negative, low and high, Fig 8B). As anticipated, WT cells overexpressing the IFNLR1-GFP chain show stronger ISG expression compared to WT cells and the magnitude of the ISG induction correlates with the relative levels of IFNLR1 expression (Fig 8B). However, the kinetic profiles of the ISGs upon type III IFN stimulation were not affected by the differential expression levels of the IFNLR1 chain (Fig 8B).

**Fig.8.**
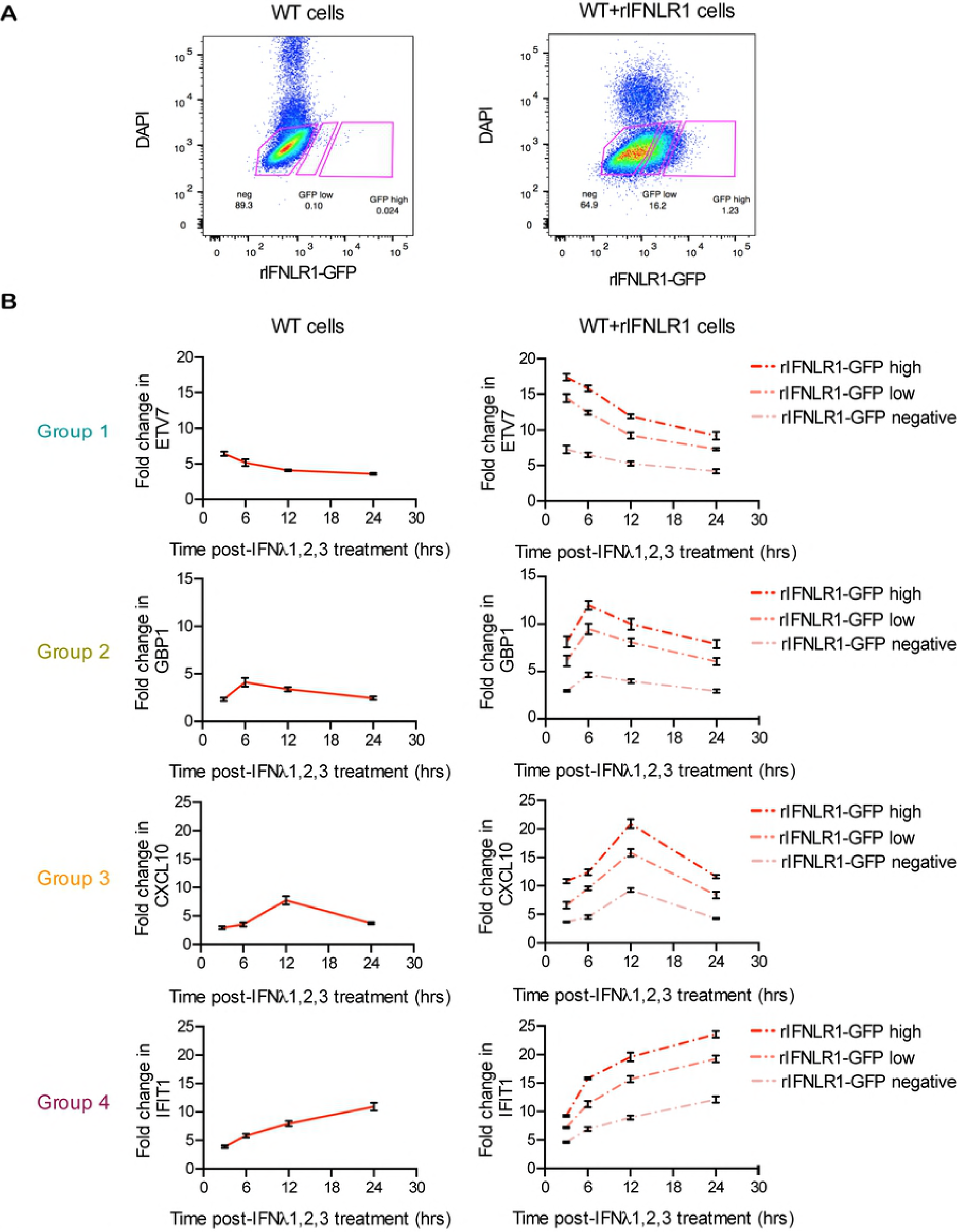
Type III IFN mediated expression kinetics of ISGs are independent of differential levels of IFNLR1 receptor. Wild-type T84 cells were transduced with rIFNLR1-GFP to create a stable line overexpressing IFNLR1 tagged with GFP. (A) WT cells overexpressing IFNLR1-GFP (WT+IFNLR1-GFP) from the same population were separated by cell sorting into three populations: non (neg)-, low-and high-expressing GFP cells. Gates were created based on the auto-fluorescence of WT cells. (B) WT and WT+IFNLR1-GFP cells were treated with type III IFN (A1-3) (300 ng/mL equivalent 13.7 nM) for 3, 6, 12 and 24 hours prior to sorting in neg-, low-and high-expressing IFNLR1-GFP cells. The kinetic pattern of expression of one representative ISG from each temporal expression patterns groups 1-4 was analyzed by qRT-PCR in each sorted population. Data are normalized to HPRT1 and are expressed relative to untreated cells at each time point. A representative experiment with technical triplicates, out of two independent experiments is shown. Mean values and SD are shown.

Altogether, our results demonstrate that type I and type III IFNs both induce an antiviral state in hIECs but with different kinetics. We could show that although both cytokines induce similar ISGs, type III IFN does it with slower kinetics and lower amplitude of individual ISG expression compared to type I IFN. Importantly, coupling mathematical modeling of both type I and type III IFN-mediated signaling and overexpression of functional IFN receptors approaches allowed us to demonstrate that these kinetic differences in type I and type III IFN ISG expression are not due to different expression level of the respective IFN receptors but are intrinsic to type I and type III IFN signaling pathways.

## Discussion

In this work, we have for the first time, performed a parallel study of the role of type I and III IFN in human mini-gut organoids and IEC lines. Our results demonstrate that type I and III IFNs are unique in their magnitude and kinetics of ISG induction. Type I IFN signaling is characterized by relatively strong expression of ISGs and confers to cells a fast-antiviral protection. On the contrary, the slow acting type III IFN mediated antiviral protection is characterized by a weak induction of ISGs in a delayed manner compared to type I IFN. Our results are in line with previous studies which also demonstrated that type III IFN is less potent than its type I IFN counterpart [5, 21, 23, 33, 34]. Additionally, we have confirmed that the delayed ISG induction seen upon type III IFN treatment of hepatocytes [21, 23, 25, 26] is not tissue specific but likely represents a global pattern of action of this cytokine in cells expressing the type III IFN receptor (i.e. human epithelial cells). In other words, the different kinetics of ISG expression induced by type I and type III IFNs are specific to each IFN signaling pathways.

In the current work, we have employed, a data-driven mathematical modeling approach to explain signal transduction kinetic differences downstream type I and type III IFN receptors. While type I IFN-mediated signaling has been previously modeled [35, 36], type III IFN has not. Our model predicted that the receptor levels directly influence the magnitude of ISG expression however, the kinetics of ISG expression appear to be intrinsic to each IFN-signaling pathway and is largely preserved under receptor overexpression. This prediction was experimentally validated by studying the response of wild-type and IFN receptor overexpressing cells to different doses of IFN (Fig 7A-C and Fig 8). This suggests that the kinetic differences in the ISG induction are intrinsic to each IFN signaling pathway. We propose that these phenotypic differences reflect functional differences, which are important for mounting a well-tailored antiviral innate immune response at mucosal surfaces where type III IFN receptors are expressed.

Both type I and III IFNs have unique and independent receptors which are structurally unrelated. These receptors are likely expressed at different levels on individual cells and their relative expression to each other might also be cell type specific. To address whether the unique ISG and antiviral expression kinetics shown by each IFN were not due to differences in their expression levels, we overexpressed into cells functional type I (rIFNAR1) and type III IFN (rIFNLR1) receptors. Our results from IFNAR1 overexpressing cells (Fig 6 and 7) are in line with previous studies showing a direct relationship between the surface levels of type I IFN receptors and the magnitude of ISG induction [37, 38]. Interestingly, we could demonstrate a similar relationship when overexpressing IFNLR1 (Fig 6 and 7) which was also associated with an increase of type III IFN antiviral potency (Sup Fig 8). These findings are in agreement with previous experiments which show that overexpression of IFNLR1 in cells which normally do not express this IFN receptor rescues both type III IFN-mediated signaling and IFN-mediated antiviral protection [5, 28]. Our IFN receptor overexpression approach demonstrates that the observed differences in ISG expression kinetics are not the results of different levels of receptors at the cell surface but is likely specific to each signal transduction pathway. Apart from the expression levels of IFN receptors, lower binding affinity towards their respective receptors could be an alternative explanation for the differential potencies of both type I and type III IFNs. Multiple studies have tried to affect the binding affinity of type I IFNs with their receptors however, results suggest that wild-type IFNs exert their antiviral activities already at maximum potency. Modifications leading to an increased affinity for their receptors do not lead to improvement of antiviral potency [32, 38-41]. To address whether the weaker activity of type III IFN could be the result of its weaker affinity for its receptor, Mendoza et al, engineered a variant of type III IFN with higher-affinity for its receptor (H11-IFNA3). They showed increased IFN signaling and antiviral activity in comparison with wild-type IFNλ3. However, the engineered variant of IFNλ3 was still acting with weaker efficacy compared to type I IFNs [32]. By exploiting the high affinity variant H11-IFNλ3, we could also show a significant increase of the amplitude of ISG expression but importantly, the kinetics of ISG expressions were not altered (Sup Fig 4).

Our results indicate a model were inherent temporal differences exist between type I and type III IFNs signaling. These differences are not the result of differential surface expression of the receptors but is the result of distinct signaling cascades from the receptors to the nucleus or within regulatory mechanisms of gene expressions.

While few studies have focused on the endocytosis and inactivation of IFNAR1, there is no information about how these processes occur for IFNLR1. It has been shown that the ternary IFNAR complex is internalized by clathrin mediated endocytosis [42] and that upon type I IFN stimulation, IFNAR1 is rapidly endocytosed and routed for lysosomal degradation [43, 44], whereas IFNAR2 can be recycled back to the cell surface or degraded [45]. Our data-driven mathematical modeling approach suggests a different kinetics of receptor activation/inactivation between both IFNs (Fig 5B and Sup Fig 5A). Therefore, further studies investigating trafficking of IFNLR1 will be important and may show that subtle changes in the time course of receptors internalization, recycling or degradation can have profound effect on kinetics of IFN activity. Apart from receptor internalization and degradation, several molecular mechanisms leading to IFN receptor inactivation have been described, such as de-phosphorylation [46, 47], or by negatively targeting the interaction of IFNAR1 with downstream signaling elements of the JAK/STAT signaling, for instance ubiquitin-specific protease USP18, and members of the suppressor of cytokine signaling protein (SOCS) family. In particular, the inhibitory role of SOCS1 in type I IFN signaling has been demonstrated in a number of previous studies, where they have shown that SOCS1 associates with TyK2 and blocks its interaction with IFNAR1 [48]. USP18 has also been shown as an important negative regulator of type I IFN signaling with a dual role acting as isopeptidase which removes the ubiquitin like-ISG15 from target proteins [49] and as a competitor of JAK1 for binding to IFNAR2 [50]. Although, limited information is available for negative regulators of the IFNLR receptor complex, the specific contribution of USP18 or SOCS in inhibition of type I versus type III IFN mediated signaling has been addressed in recent studies. In particular, it has been showed that both type I and III IFNs (IFNa, IFNp and IFNA1, A2, A3 and A4) induced the expression of USP18, SOCS1 and SOCS3 [51-57] and overexpression of all these negative regulators inhibited both IFNa and IFNA1 mediated JAK-STAT signaling [54, 56] suggesting that at ‘’supraphysiological’’ expression levels all the inhibitors can block type I and type III mediated JAK-STAT signaling [56]. Additionally, it has been shown that USP18 is induced later and that its level increased over time, correlating with the long lasting refractories of IFNa signaling [51, 52, 56]. In our study we observed also a later peak of induction of USP18 at 12h or 24h upon type I or type III IFNs, respectively. In line with the above-mentioned studies we also observed rapid and transient induction of SOCS1 upon type I IFN treatment and sustained induction upon type III IFN stimulation. However, further investigation is required to determine the correlation of the kinetics of induction of these negative regulators with the ISGs induction in type I versus type III IFN treatment in human IECs.

In the canonical type I and III IFN signaling pathway the next downstream players from the IFN receptors are the JAKs, STAT1, STAT2 and IRF9, which are all regulated on the level of expression and activation. Our own observations (data not shown) and previous studies could not explain the major differences in the kinetics of type I versus type III IFNs activity by focusing on the time course of phosphorylation of STATs [21, 25]. However, given that alternative modifications of STATs *(e.g.* phosphorylation on alternative residues, acetylation, methylation and sumoylation patterns) have been proposed to contribute to the activity of type I IFNs [26, 58-60] it might be possible that new modifiers of STAT activity may determine the kinetic pattern of action of type I versus type III IFNs. In addition, apart from the JAK/STAT axis, there is accumulating evidence which correlates ISG transcription upon IFN treatment with a plethora of JAK-STAT independent pathways, such as members of the CRK [61-63] and MAPKinase family [24, 28, 64-66], which might also temporally coordinate IFNs kinetic profile of action. Apart from the differences in the signaling cascade of type I versus type III IFNs, an explanation for their differential kinetics of action might stem from the physiology of the different cell types. For example, in a recent study Bhushal et al. reported that polarization of mouse intestinal epithelial cells eliminates the kinetic differences between type I and type III IFNs, by accelerating type III IFN responses [33, 67].

Several studies describing the transcriptional activities of both type I and type III IFNs have reported that very similar sets of ISGs are produced upon both type I and III IFN stimulation [12, 17, 21, 22, 25, 28] while only few ISGs appear to be predominantly expressed upon type III IFN treatment in murine IECs [67]. We believe that there are several functional advantages for adopting a lower and slower activity, like the profile of action of type III IFN, in the antiviral protection of epithelial tissues. The differences in the temporal expression of ISGs could create unique antiviral environments for each IFN. Many ISGs function as pro-inflammatory factors [30, 68]. By stimulating ISGs production in high magnitude, an excessive amount of antiviral and proinflammatory signals could be produced which on the one hand will eliminate efficiently viral spreading but on the other hand may cause local exacerbated inflammation and irreversible tissue damage, leading to chronic inflammation in mucosal surfaces.

In addition, the expression of different functional groups of ISGs at early and at late time points (Fig 3) might allow cells to create two distinct phases within the antiviral response. At early time points, minimum levels of ISGs may act to protect the host against viral infection. Antiviral ISGs will be responsible for fighting the pathogens and pro-inflammatory ISGs will stimulate members of the adaptive immune system to assist the antiviral protection. At later time points the produced ISGs, may be involved in antiinflammatory processes, such as resolving of inflammation and tissue healing and repair [66, 69]. To exert this anti-inflammatory role, ISGs may need to be produced in higher levels, as they might act more paracrine and spread through the tissue to balance again the tissue homeostasis after the viral attack. In conclusion, we propose that type III IFN-mediated signaling is not only set to act predominantly at epithelium surfaces due to the restriction of its receptor but also is specifically tailored to mount a distinct immune state compared to other IFNs which is critical for mucosal surfaces that face the challenge.

## Materials and Methods

### Antibodies/Reagents

Commercially available primary antibodies were mouse monoclonal antibodies recognizing beta-Actin (Sigma #A5441), phospho STAT1 and STAT1 (BD Transductions #612233 and #610115, respectively). Anti-mouse (GE Healthcare #NA934V), coupled with horseradish peroxidase was used as secondary antibody for Western blot at a 1:5000 dilution. Human recombinant IFN-beta1a (IFNβ) was obtained from Biomol (#86421). Recombinant human IFNλ1 (IL-29) (#300-02L) and IFNλ2 (IL28A) (#300-2K) were purchased from Peprotech and IFNλ3 (IL-28B) from Cell signaling (#8796). High affinity engineered IFNA3 variant (H11) and wild type IFNλ3 were produced as described in [32]. The IFN concentrations used to treat the cells are stated in the main text and in the figure legends.

### Cell and Viruses

T84 human colon carcinoma cells (ATCC CCL-248) were maintained in a 50:50 mixture of Dulbecco’s modified Eagle’s medium (DMEM) and F12 (GibCo) supplemented with 10% fetal bovine serum and 1% penicillin/streptomycin (GibCo). SKCO15 cells were maintained in DMEM with 10% fetal bovine serum, 1% penicillin/streptomycin, 15mM HEPES and 1% NEAA (Non-Essential Amino Acids). Mini-gut organoids were harvested and maintained as described earlier [24]. VSV-Luc was used as previously described [24].

### Ethics Statement

Human colon tissue was received from colon and small intestine resection from the University Hospital Heidelberg. This study was carried out in accordance with the recommendations of “University Hospital Heidelberg” with written informed consent from all subjects. All subjects gave written informed consent in accordance with the Declaration of Helsinki. All samples were received and maintained in an anonymized manner. The protocol was approved by the “Ethic commission of University Hospital Heidelberg” under the approved study protocol S-443/2017.

### RNA isolation, cDNA, and qPCR

RNA was harvested from cells using NuceloSpin RNA extraction kit (Macherey-Nagel) as per manufacturer’s instructions. cDNA was made using iSCRIPT reverse transcriptase (BioRad) from 200ng of total RNA as per manufacturer’s instructions. qRT-PCR was performed using SsoAdvanced SYBR green (BioRad) as per manufacturer’s instructions, TBP and HPRT1 were used as normalizing genes.

### Gene expression analysis of interferon stimulating genes

Colon organoids and T84 cells were treated with 2000 RU/ml of type I IFN (β) or 100 ng/ml of each type III IFN (λ1,2 and 3). Total RNA was isolated at 3, 6, 12 and 24h post-treatment as described above. For the gene expression analysis of interferon stimulated genes (ISGs), qRT-PCR was performed using the predesigned 384-well assay of type I IFN response for use with SYBR Green assaying the expression of 87 ISGs (Biorad # 10034592). The expression of 45 additional ISGs and transcriptional factors was analyzed by qRT-PCR with primer sets obtained as previously described [27]. The complete gene list monitored in this study and the primers used to amplify each gene is available in Tables S1 and S2. Differential expression analysis of each treatment was performed by comparing the baseline expression of genes in an untreated control at each time point. Only genes which were either induced or reduced more than 2-fold in any of the samples were considered to be significantly regulated. These genes were either analyzed using scatterplots or visualized by a heatmap after sorting the fold change of expression in response to type I IFN (β) in decreasing order. For the T84 cells all fold change values above 20 and below 0.05 were replaced with 20 and 0.05 respectively. For the organoids, the fold change values above 800 and below 1/800 were replaced with 800 and 1/800. This data adaptation was done to center the heatmap around 0 (white) and to avoid errors in logarithmic calculations. When visualizing the expression peaks, only the highest value is shown per time point for each gene. All analyses were performed using R version 3.3.0 and 3.3.3 including the packages gplots and ggplot2.

### Western blot

At time of harvest, media was removed, cells were rinsed one time with 1X PBS and lysed with 1X RIPA buffer (150 mM sodium chloride, 1.0% Triton X-100, 0.5% sodium deoxycholate, 0.1% sodium dodecyl sulphate (SDS), 50 mM Tris, pH 8.0 with phosphatase and protease inhibitors (Sigma-Aldrich)) for 20mins at 4°C. Lysates were collected and equal protein amounts were separated by SDS-PAGE and blotted onto a PVDF membrane by wet-blotting. Membranes were blocked with 5% milk or 5% BSA, when the phospho STAT1 antibody is used, in TBS containing 0.1% Tween 20 (TBS-T) for one hour at room temperature. Primary antibodies were diluted in blocking buffer and incubated overnight at 4°C. Membranes were washed 4X in TBS-T for 15mins at RT. Secondary antibodies were diluted in blocking buffer and incubated at RT for 1h with rocking. Membranes were washed 4X in TBS-T for 15mins at RT. HRP detection reagent (GE Healthcare) was mixed 1:1 and incubated at RT for 5mins. Membranes were exposed to film and developed.

### VSV luciferase assay

Colon organoids and T84 cells were seeded in a white F-bottom 96-well plate. Samples were pre-treated prior to infection or treated post-infection as indicated with increasing concentrations of type I or type III IFNs. VSV-Luc was added to the wells and the infection was allowed to proceed for 8hrs. At the end of the infection, media was removed, samples were washed 1X with PBS and lysed with Cell Lysis Buffer (Promega) at RT for 20 mins. A 1:1 dilution of Steady Glo (Promega) and Lysis Buffer were added to the samples and incubated at RT for 15 mins. Luminescence was read using an Omega Luminometer.

### FACS analysis

Fluorescence-activated cell sorting (FACS) was performed on FACSMelody™ Cell Sorter (BD Biosciences). DAPI was added for nuclear staining. Data were processed using FlowJo 10.0.5.

### Cloning and generation of stable cell lines

Knockout of IFNAR1 and IFNLR1 in T84 cells were achieved by using the CRISPR/Cas9 system as described earlier [24]. For back-compensation of the IFN receptor KO cell lines and for generation of wild-type T84 cells overexpressing the IFNAR1 and IFNLR1, plasmids containing the cDNA of IFNAR1 and IFNLR1 were obtained from a gateway compatible ORF bank (pENTRY221-IFNAR1) and from GE Healthcare (pCR_XL_TOPO_IFNLR1, #MHS6278-213246004), respectively. The IFNLR1-GFP construct (pC1-HsIFNLR1-GFP) was generated using the following cloning strategy. A mammalian expression plasmid producing a N-terminal EGFP-tagged extracellular domain of IFNLR1 (EGFP-IFNLR1) was generated as follows: cDNA corresponding to this open reading from was generated synthetically (GeneArt, Life Technologies) and subsequently sub-cloned directly into the pC1 expression plasmid (Promega) backbone. Specifically, monomeric EGFP was introduced between the signal peptide sequence and the remaining glycoprotein flanked by three alanine residues at its amino terminus and a short glycine-serine linker sequence of N-AAASGSGS-C at its carboxyl terminus. Tri-alanine flanking allowed facile incorporation of restriction enzyme sites (Not1 and SacII) allowing removal or swapping of EGFP tag. Sequences available on request. Caspase-cleavage resistant IFNAR1 and IFNLR1 were generated using the Quick Change II XL site directed mutagenesis kit (Agilent Technologies, Germany), following manufacturer’s instructions. Point mutations were controlled by plasmid sequencing. The expression vectors were generated by inserting the respective constructs into the lentiviral vector pDest GW35 by using the Gateway cloning technology (Life Technologies, Germany) according to manufacturer’s instructions. Lentiviruses were produced as previously described [24], and T84 cells were transduced two times using concentrated stocks of lentiviral particles encoding the cleavage resistant IFNAR1 and IFNLR1. 36 hours posttransduction, transduced cells were selected for using blasticidin.

### Model simulation and parameter estimation

The mathematical model was implemented in terms of ordinary differential equations (ODEs) in MATLAB 2016b (S3 Table). The numerical simulations were conducted using the CVODES, a module from SUNDIALS numerical simulation package, in the MATLB environment. The model was initially set to a steady state condition and most of the initial conditions were set (S4 Table). Only, the IFNLR efficacy factor was estimated using time-resolved ISG expression data that we measured with different doses of type I IFN (β) or III IFN (λ1-3). All of the ISG expression data for the IFNAR1 and IFNLR1 overexpression experiments were reproduced only by fitting new initial values of IFNAR1 and IFNLR1 (S5 Table).

Parameter estimation was conducted by minimizing the weighted nonlinear least squares,

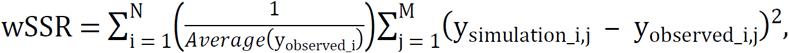
of model simulations versus data points, j = 1, …, M, of different experiments, i = 1, …, N. The inverse of the average of every time-resolved experimental data was used as a weighting factor for fitting the corresponding data.

### Profile-likelihood analysis

To assess the uncertainty in the estimated parameter values, we used the profile-likelihood method [70]. In this method, the parameter confidence bounds are calculated based on their contribution to the likelihoods, or in another word, the objective function (wSSR). This computational approach is conducted in a stepwise manner. In every step, the respective parameter is fixed at a new value distant from the optimum estimated one. Then, the new maximum likelihood is calculated (wSSR_min_(0)). Using this approach, we can calculate the profile of the maximum likelihoods over different values of the considered parameter. Then a threshold, Δ_a_,

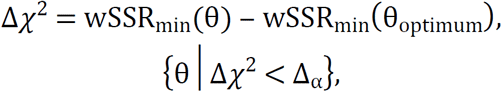
is used to define the confidence bounds for the respective parameter. The threshold, Δ_α_, is the α quantile of the chi-squared distribution.

### Approximate 95% confidence bands calculation

To investigate the effect of the parameter uncertainty on model predictions we calculated approximate 95% confidence bands, as explained in Seber and Wild [71].

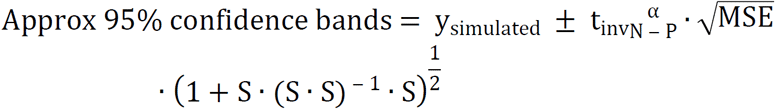
where “t_invN-P_^α^” is the a quantile of student’s t distribution, “N” is the number of data points and “P” is the number of estimated model parameters, “MSE” is the mean standard error and “S” is the sensitivity matrix of the respective simulated observable.

### Model selection

To select the most parsimonious model, the simplest model with good predictive power, from the ensemble of the four alternative models of the ISG response to type I versus type III interferon, we used the Akaike information criterion corrected for small sample size (AICc). After fitting the models to the experimental data, we calculate the AICc score for every model. AICc is calculated as:

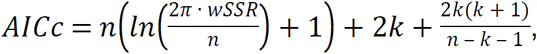
where n is the number of data points used to fit the model, k is the number of estimated parameters of the respective model, and wSSR is the minimum weighted sum of squared residuals for the respective model. The model with the minimum AICc value is selected as the most parsimonious model from the ensemble of alternative models. In order to compare the selected model with other models, we calculate both ΔAICc, the difference between the AICc value of the models with the minimum AICc value from the ensemble of the models, and the AICc weight (w i). The Akaike weight is a weight of evidence for the respective model and is calculated as:

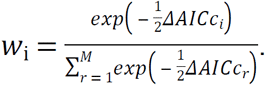

## Acknowledgments

We are grateful to the following colleagues for providing reagents Ronald L. Rabin and Lynnsey A. Renn (Center for Biologics Evaluation and Research, US Food and Drug Administration, USA) for the transcription factor primers, Sean Whelan (Harvard Medical School, USA) for the VSV-Luc and Asma Nusrat (University of Michigan, USA) for the SKCO15 cells. We would like to thank the members of Steeve Boulant Lab (Heidelberg University Hospital, Germany) for their insights and helpful discussions in preparing the manuscript. We would like to thank the FACS facility at the EMBL, Heidelberg for their assistance in sorting the INFLR-GFP expressing cells.

## Supporting Information Legends

**Sup Fig 1. Kinetics of type I and type III IFN-mediated antiviral activities in intestinal organoids.** (A-B) Intestinal organoids were pre-treated with the indicated concentrations of type I IFN (β) or type III IFN (λ1-3) for 2.5 h prior to infection with VSV-Luc using a multiplicity of infection (MOI) of 1. Viral replication was assayed by measuring the luciferase activity. (A) The relative antiviral protection is expressed as a percentage of total protection in VSV-infected organoids or (B) as the EC90 corresponding to the concentration of type I IFN (β) or type III IFN (λ1-3) resulting in 90% inhibition (10% infection) of viral replication. (C-D) Intestinal organoids were treated with type I IFN (β) (2,000 RU/mL equivalent 0.33 nM) or type III IFN (λ1-3) (100ng/mL each or total 300 ng/mL equivalent 13.7 nM) for different times prior to infection with VSV-Luc. Viral replication was assayed by measuring luciferase activity. (C) The relative VSV infection is expressed as the percentage of the luciferase activity present in VSV-infected organoids without IFN treatment (set to 100). (D) Pre-incubation time of type I IFN (β) or type III IFN (λ1-3) required to inhibit VSV infection to 10% (90% inhibition). (E-F) Same as (C-D), except intestinal organoids were treated at the indicated times post-infection with VSV-Luc. (F) Delayed-time post-infection for type I IFN (β) or type III IFN (λ1-3) to still inhibit VSV infection to 90% (10% inhibition). Data represent the mean values of two independent experiments with intestinal organoids generated from two different donors. Error bars indicate the SD. *<P.05, **P < 0.01, ***P < 0.001, ns, not significant (unpaired t-test).

**Sup Fig 2. Kinetics of type I and type III IFN-mediated antiviral activities in human intestinal epithelial cells.** (A-B) T84 cells were pre-treated with the indicated concentrations of type I IFN (β) or type III IFN (λ1-3) for 2.5 h prior to infection with vesicular stomatitis virus (VSV) expressing Firefly luciferase (VSV-Luc) using a multiplicity of infection (MOI) of 1. Viral replication was assayed by measuring the luciferase activity. (A) The relative antiviral protection is expressed as a percentage of total protection in VSV-infected cells or (B) as the EC90 corresponding to the concentration of type I IFN (β) or type III IFN (λ1-3) resulting in 90% inhibition (10% infection) of viral replication. (C-D) T84 cells were treated with type I IFN (β) (2,000 RU/mL equivalent 0.33 nM) or type III IFN (λ1-3) (100ng/mL each or total 300 ng/mL equivalent 13.7 nM) for different times prior to infection with VSV-Luc. Viral replication was assayed by measuring luciferase activity. (C) The relative VSV infection is expressed as the percentage of the luciferase activity present in VSV-infected cells without IFN treatment (set to 100). (D) Pre-incubation time of type I IFN (β) or type III IFN (λ1-3) required to inhibit VSV infection to 10% (90% inhibition). (E-F) Same as (C-D), except T84 cells were treated at the indicated times post-infection with VSV-Luc. (F) Delayed-time post-infection for type I IFN (β) or type III IFN (λ1-3) to still inhibit VSV infection to 90% (10% inhibition). Data in (A-F) represent the mean values of three independent experiments. Error bars indicate the SD. *<P.05, **P < 0.01, ***P < 0.001, ****P < 0.0001, ns, not significant (unpaired t-test).

**Sup Fig 3. Type III IFNs have a lower transcriptional activity compared to type I IFNs in human intestinal epithelial cells.** (A-B) T84 cells were stimulated with indicated concentrations of type I (β) or III IFN (λ1-3) for different times and the transcript levels of the ISGs IFIT1 and Viperin were analyzed by qRT-PCR. Data are normalized to TBP and HPRT1 and are expressed relative to untreated cells at each time point. A representative experiment with technical triplicates, out of three independent experiments is shown. Mean values and SD are shown. (C-D) T84 cells were treated with type I IFN (β) (2,000 RU/mL equivalent 0.33 nM) or type III IFN (λ1-3) (300 ng/mL equivalent 13.7 nM) for the indicated times and identification of the IFN-induced ISGs was performed by qRT-PCR. A total of 70 out of 132 ISGs tested were found to be significantly induced more than 2-fold compared with a baseline (mean of untreated controls at the particular time points) for at least one time point by at least one IFN treatment. Data are normalized to TBP and HPRT1. (C) Comparison of expression values (log2 (Fold Change)) for all genes induced at the indicated times with type I IFN (β) versus type III IFN (λ1-3). Solid line indicates equivalent expression. (D) Hierarchical clustering analysis of these genes produced four distinct temporal expression patterns (Groups 1-4) based on the time-point of the maximum induction in response to type I IFN (β) or type III IFN (λ1-3). Color codes have been used to visualize the induction peak per group. Gene expression heat map showing the genes clustered in their respective temporal expression patterns groups in response to type I IFN (β) or type III IFN (λ1-3). The genes per group are sorted in decreasing order on the basis of their fold change of expression in response to type I IFN (β) or type III IFN (λ1-3) and only showing the highest expressed values within the temporal groups omitting all other values for visualization.

**Sup Fig 4. Comparison of the transcriptional response between wild-type IFNA3 and high affinity H11-IFNλ3 variant.** (A-D) T84 cells were stimulated with increasing concentrations of type III IFN (λ3) (WT-IFNλ3) or the high affinity IFNλ3 variant (H11-IFNλ3) for indicated times and the kinetic of expression of one representative ISG from each temporal expression groups 1-4 was analyzed by qRT-PCR, (left column) WT-IFNλ3, (right column) H11-IFNλ3 treated cells. Data are normalized to HPRT1 and are expressed relative to untreated cells at each time point. A representative experiment with technical triplicates. Mean values and SD are shown.

**Sup Fig 5**. **Analysis of mathematical model M3**. (A) Comparative simulation of type I and type III IFN receptor complex activation. Cellular concentration of the activated type I or type III IFN receptor complex, upon treatment with 0.1 nM of IFNs, are simulated using the calibrated model. (B) Profile likelihoods of model parameters. The uncertainty of the estimated model parameters is calculated using the profile likelihood method. The solid blue line is the change in the weighted sum of squared residuals (Ax^2^), the filled circle indicates the optimum parameter value, and the solid red line indicates the 95% threshold calculated using the x2 distribution. (C) The 95% confidence bounds of type I or type III IFN receptor complex inactivation rate constants are calculated using the profile likelihood method. Our calculations show that the type III IFN receptor complex inactivation rate constant (k2) is significantly smaller than the corresponding rate constant for type I IFN receptor complex (k11). (D-E) The mathematical model shows the effect of IFNAR1 (D) and IFNLR1 (E) overexpression of up to 3-fold (3*IFNAR1 and 3*IFNLR1) on Viperin activation upon treatment with representative concentrations of type I IFN (β) (0.1 nM) or type III IFN (λ1-3) (13.7 nM), respectively.

**Sup Fig 6. Expression levels of IFN receptors in T84 cells.** (A) T84 wild-type cells, (B) T84 cells overexpressing rIFNAR1 (WT+rIFNAR1) and (C) T84 cells overexpressing rIFNLRI (WT+rIFNLR1) were analyzed by qRT-PCR to quantify the transcript levels of IFNAR1, IFNAR2, IFNLR1 and IL10RB (IFNLR2). Data are normalized to HPRT1. The mean value obtained from three independent experiments is shown. Error bars indicate the SD. *<P.05, **P < 0.01, ***P < 0.001, ****P < 0.0001, ns, not significant (unpaired t-test).

**Sup Fig 7. Type I and type III IFN receptors are functional when overexpressed into cells.** (A-D) T84 IFNAR1-/-cells were rescued by stable expression of a cleavage resistant mutant of rIFNAR1 (see methods for details) (T84 IFNAR1-/-+ rIFNAR1). (B-D) T84 IFNAR1-/-, T84 IFNAR1-/-+ rIFNAR1 cells and control T84 cells scramble gRNA (SCR) were pre-treated with type I IFN (β) (2,000 RU/mL equivalent 0.33 nM) or type III IFN (λ1-3) (300 ng/mL equivalent 13.7 nM). (B) 2.5 h post-treatment, T84 cells were infected with VSV-Luc (MOI = 1). Viral replication was assayed by measuring the luciferase activity. For each sample luciferase activity was measured in triplicates and is expressed as the percentage of the luciferase signal in VSV-infected cells without IFN treatment (set to 100) for each cell lines. (C) 1h post IFN treatment, IFN signaling was measured by immunoblotting for pSTAT1 Y701. Actin was used as a loading control. A representative immunoblot out of three independent experiments is shown. (D) same as (B), except that induction of IFN-stimulated genes was monitored by relative qRT-PCR quantification of IFIT1 and Viperin at the indicated times post-IFN treatment. (E-H) same as (A-D) except that T84 IFNLR1-/-were rescued by stable expression of a cleavage resistant mutant of rIFNLR1 (T84 IFNLR1-/-+ rIFNLR1). Data were normalized to HPRT1 and are expressed relative to untreated cells of each time point. The mean value obtained from three independent experiments is shown. Error bars indicate the SD.

**Sup Fig 8. Establishment of an antiviral state in cells overexpressing the IFN receptors and treated with IFNs.** (A-H) Wild-type T84 cells were transduced with rIFNAR1 or rIFNLR1 to create stable lines overexpressing either receptor. (A-D) T84 wild-type cells (WT) and T84 cells overexpressing the IFNAR1 (WT+rIFNAR1) were treated with (A-B) type I IFN (β) (2,000 RU/mL equivalent 0.33 nM) or (C-D) type III IFN (λ1-3) (100ng/mL each =300 ng/mL equivalent 13.7 nM) at the indicated times (A, C) prior to infection or (B, D) post infection with VSV-Luc. Viral replication was assayed by measuring the luciferase activity. The time necessary to confer IECs an antiviral state was addressed by measuring the impact of IFN treatment on viral replication. For each sample luciferase activity was measured in triplicates and is expressed relative to VSV-infected cells without IFN treatment (set to 100). (E-H) same as (A-D), but T84 wild-type cells (WT) and T84 cells overexpressing the IFNLR1 (WT+rIFNLR1) were treated with (E-F) type I IFN (β) (2,000 RU/mL equivalent 0.33 nM) or (G-H) type III IFN (λ1-3) (100ng/mL each =300 ng/mL equivalent 13.7 nM) at the indicated times (E, G) prior to infection or (F, H) post infection with VSV-Luc. Data represent the mean values of three independent experiments. Error bars indicate the SD. *<P.05, **P < 0.01, ns, not significant (unpaired t-test).

**Sup Fig 9. Expression kinetics of ISGs are independent of the IFN receptor levels.** (A-B) Wild-type T84 cells were transduced with rIFNAR1 or rIFNLR1 to create stable lines overexpressing either receptors. (A) T84 wild-type cells (WT) and T84 cells overexpressing the IFNAR1 (WT+rIFNAR1) were treated with increasing concentrations of type I IFN (β) for the indicated times and the expression kinetics of the ISG VIPERIN were analyzed by qRT-PCR. Data are normalized to HPRT1 and are expressed relative to untreated cells at each time point. (B) Same as (A), except T84 cells overexpressing the IFNLR1 (WT+IFNLR1) were used and treated with increasing concentrations of type III IFN (λ1-3). A representative experiment with technical triplicates, out of three independent experiments is shown. Mean values and SD are shown. (C-D) The mathematical model predicts the effect of IFNAR1 and IFNLR1 overexpression on Viperin activation upon treatment with different concentrations of type I IFN (β) and type III IFN (λ1-3). The IFNAR1 and IFNLR1 levels were increased ~2.6 and ~1.5 fold, respectively, while all other parameters were held constant. The solid lines are the best fits and the shaded areas are 95% confidence intervals. (D) The mathematical model correctly predicts the IFNAR1 versus IFNLR1 overexpression, measured experimentally by qRT-PCR.

**Sup Table 1. List of primers used in predesigned 384 well assay qRT-PCR.** For the gene expression analysis of interferon stimulated genes (ISGs), qRT-PCR was performed using a predesigned 384-well assay of type I IFN response assaying the expression of ISGs. The Reference Sequence (RefSeq) accession number is provided for each ISG tested.

**Sup Table 2. List of primer sequences used for qRT-PCR analysis.** The expression of additional ISGs, transcriptional factors and housekeeping genes was analyzed by qRT-PCR with the primer sets shown in this table.

**Sup Table 3. Mathematical formulation of the model.** The table lists all differential equations explaining the dynamics of different biological species in our model. Cell surface (Area) is calculated as, Area = (36n)^1/3^V_cell_^2/3^. Cell volume (V_cell_) is assumed equal to 2*10^-9^ Liter. Brackets [ ] indicate the concentration of the respective biological species.

**Sup Table 4. State variables and initial values.** Biological species considered in the model (state variables) and their initial values are listed in the table.

**Sup Table 5. Estimated parameter values.** The estimated value of the model free parameters, their profile-likelihood based confidence bound and their dimensions are explained in the table. All reaction rate constants of the model, k_1_-k_9_, are practically identifiable.

